# Sidekick2 facilitates multiciliated cell penetration through multicellular adherens junctions

**DOI:** 10.64898/2026.06.03.729971

**Authors:** Eve E. Suva, Tomohito Higashi, Brian J. Mitchell

## Abstract

Multicellular vertices are important cellular interfaces for mediating tissue dynamics across the epithelium. Here, we report that expression of the vertebrate adhesion protein, Sidekick2 (Sdk2), localizes to vertebrate multicellular adherens junctions in the embryonic *Xenopus laevis* epithelium. Using the developmental process of radial intercalation, we find Sdk2 facilitates multiciliated cell penetrance into the outer epithelium at multicellular junctions. With fluorescence recovery after photobleaching, we characterize Sdk2 dynamics at multicellular and bicellular junctions and interrogate how the intracellular and extracellular domains of Sdk2 contribute to these dynamics. We observe that overexpression of the highly conserved C terminal intracellular domain alone but not the extracellular domain alone is sufficient to act as a dominant negative, delaying radial intercalation. These results implicate Sdk2s intracellular interactions as critical for regulating the junctional remodeling required during radial intercalation. Ultimately, our findings implicate Sdk2 as a component of multicellular vertices that aids the governance of epithelial integrity.

## Introduction

Preservation of epithelial barrier function is essential in maintaining homeostasis. As the primary shield protecting multicellular organisms, epithelia must dynamically adjust to both mechanical and chemical cues in the environment. A subtype of epithelial adjustment, intercalation, describes the process of neighboring cells exchanging places and is crucial in embryonic development^1^. Intercalation is mediated by remodeling the junctional proteins that connect individual epithelial cells into a tissue. While much is known about mediolateral intercalation (MLI)^1–4^, neighboring cells in a single plane exchanging places, less is understood about the process of radial intercalation (RI)^5^, cells from apical or basal layers passing through a multicellular tissue. Understanding the junctional interactions during RI is crucial for understanding normal developmental processes and could provide insights into diseases associated with defects in epithelial integrity, such as diseases involving chronic inflammation^6^ or cancer metastasis^7^.

Vertebrate epithelial junctions are composed of apically positioned tight junctions (TJ) that seal the exterior, and sup-apical Adherens junctions (AJ) and desmosomes, which fasten cells together in response to mechanical cues. Taken together, these layers of junctional proteins form bicellular junctions (BcJ), binding two epithelial cells together along their shared sides. In contrast, specialized junctions occur at multicellular junctions (McJs), which are vertices where three or more cells connect. McJs are crucial to epithelial integrity as they are recognized as important sites of tension and cellular transport^8^. Numerous studies of vertebrate epithelium have identified multicellular tight junction (McTJ) proteins, namely Tricellulin^9^ (Tric) and the LSR^10–12^ family proteins. In contrast, the role of AJ proteins at vertebrate McJs remains poorly characterized despite recent advances from *Drosophila* that implicate Sidekick (Sdk) as an important AJ protein enriched at McJ^13–15^.

First identified as a component of Drosophila eye development^16^, Sdk has now been shown to localize to multicellular adherens junctions (McAJ) throughout the epithelium of embryonic *Drosophila*^13–15^. While invertebrates have one Sdk, vertebrates typically have two, Sdk1 and Sdk2^17^. Both Sdk1 and Sdk2 were identified in retinal studies as important for vertebrate eye development^18^. Expression of Sdk1 and Sdk2 has been observed broadly during embryonic development in mice^19^; however, most vertebrate studies have focused on retinal and neuronal tissues^18,20–23^.

As one of the largest members of the immunoglobulin (Ig) superfamily, Sdk proteins have six Ig domains on the N-terminus followed by thirteen fibronectin type III (FNIII) repeats, then a single-pass transmembrane domain with a relatively short intracellular domain ending in a highly conserved PDZ binding motif (-SXV) at the C-terminus^17^. Previous X-ray crystallography studies with mouse (Ms) Sdk1 and Sdk2 suggest the first four Ig domains form a curved ‘horseshoe’ structure to dimerize, with the first and third Ig domains of one Sdk binding with the second and fourth of an opposing Sdk^24^. Additional in vitro studies suggest the 13 FNIII repeats lie along the descending sides of the cell membrane at the junctions^25^, which would account for the basolateral string-like structures observed with Sdk in *Drosophila* epithelium^14^. Compared with the extracellular domain, the relatively short intracellular region of Sdk proteins is intrinsically disordered and terminates in a PDZ-binding domain^17^. Prior assays in Drosophila have shown interactions with other junction-associated proteins such as ZO-1 (Pyd^13,26^ in Drosophila), Afadin (Cno^13,26^ in Drosophila), and the WAVE regulatory complex^27^ at this PDZ binding domain. Both the extracellular and intracellular domains of Sdk have roles in junctional remodeling during *Drosophila* development, as seen with MLI T-1 transitions^13–15,27^, and with RI during trachea development^13,26^.

In this study, we use the embryonic *Xenopus laevis* epithelium to address how Sdk2 contributes to epithelial integrity. We studied its role in the context of RI of multiciliated cells (MCCs), which are known to penetrate the epithelium specifically at McJs^28–30^. We observe that Sdk2 impacts the timing of MCC RI. Furthermore, we find disruptions to the extracellular or intracellular domains alter Sdk2 dynamics and observe that removal of the extracellular domain delays MCC RI.

## Results

### Sdk2 Localizes to Vertebrate Adherens Junction Vertices

Sdk2 is expressed during *Xenopus* embryonic development^31^. To examine the localization of Sdk2 in vertebrate epithelium, we injected *Xenopus laevis* embryos with mRNA encoding the Ms copy of the Sdk2 gene with a GFP tag on the C-terminus, Sdk2-GFP (Figure 1A). We observed Sdk2-GFP along the junctions of *Xenopus* epithelium, with a substantial enrichment at McJs (Figure 1B), similar to observations in *Drosophila*^14^. This increased intensity at the vertices notably descends basolaterally along junctions, suggestive of localization below the TJs (Figure 1C).

**Figure 1:**
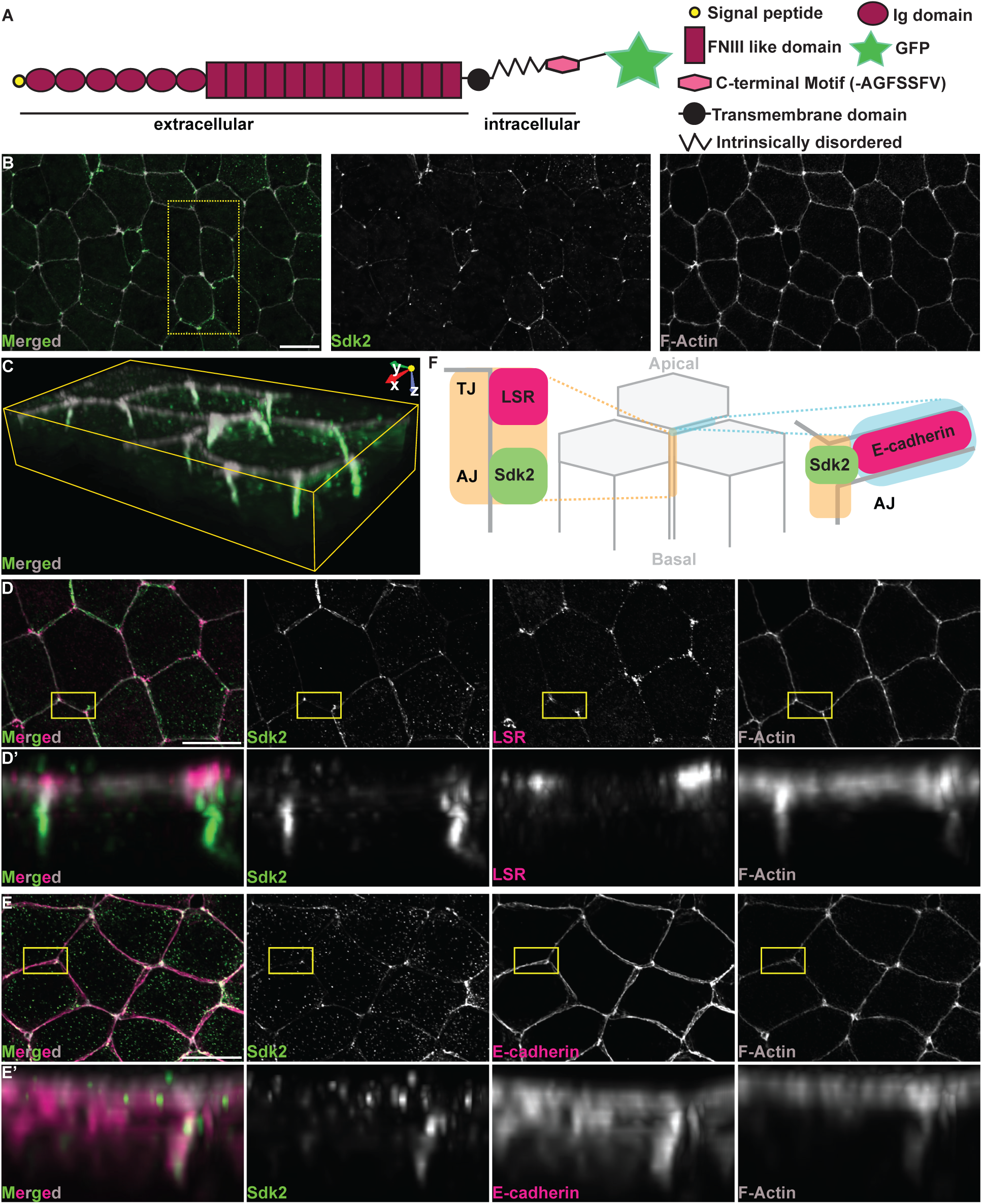
Vertebrate Sidekick2 is an adherens junction protein enriched at multicellular vertices. A) Cartoon depiction of the domain architecture for Sdk2-GFP. (B-C) Z projections of ST22 embryo marked with MsSdk2 GFP (green) F-Actin (gray) with the outlined area 3D tilted in (C). (D-E) Z projections of a ST21 embryo marked with MsSdk2-GFP (green) F-Actin (gray) together with LSR antibody (pink) (D) and Ecandherin-2xMcherry (pink) (E). (D’-E’) Side projections from boxed regions in (D and E). (F) Cartoon depiction of Sdk2 localization at McAJ. Orange shading represents the McJ and Light blue shading represents a BcJ. (A, D, E) Scale bars are 20µm.

To determine the junctional placement of Sdk2, we compared its localization with known TJ and AJ proteins. We observed Sdk2-GFP below the McTJ proteins LSR (Figure 1D) and Tric-RFP (Figure S1A). Even the Sdk2-GFP observed at BcJ is subapical, as shown when co-expressed with the general TJ protein Occludin-mCherry (Figure S1B). This indicates that Sdk2 is not a TJ protein but instead localizes at the level of AJs of vertebrate epithelium. To confirm this, we examined Sdk2 localization in comparison with the well-known AJ protein, E-cadherin, and observed colocalization of Sdk2-GFP with either co-expression of E-cadherin-2xmcherry (Figure 1E) or antibody staining of endogenous *Xenopus* E-cadherin (Figure S1C). These findings indicate that Sdk2 is a McAJ protein in the vertebrate epithelium (Figure 1F).

### Sdk2 influences the timing of Multiciliated Cell Radial Intercalation

We next explored the role of Sdk2 in maintaining epithelial integrity. Within the epithelium of *Xenopus* embryos, MCCs are initially born in a sublayer of the epithelium and undergo a developmentally timed RI, whereby they insert into the outer epithelium specifically at McJs^28,30,32^. Importantly, we have previously shown that the timing of RI is both positively and negatively malleable, providing a quantifiable assay for the efficiency of insertion^33,34^. By scoring the total number of MCCs, compared with MCCs that have successfully penetrated the outer epithelium and have achieved an apical surface area of ≥30*μ*m^2^, we can quantify the efficiency of RI events at a given time in development. We first compared the effects of overexpression (OE) of Sdk2 on MCC RI. We injected only one cell of a two-cell embryo, resulting in a mosaic embryo where only half the embryo expresses the injected mRNA, leaving the other side as an uninjected control (Figure 2A). Upon mosaic injection, we observed more successful MCC insertions at stage (ST) 20 (i.e. apical surface of 30*μ*m^2^ ≥, denoted with white arrowheads); whereas, the uninjected control had fewer successful insertions (30*μ*m^2^ ≤, denoted with yellow arrows) (Figure 2B-C). Quantification revealed a significant increase of MCC insertion with Sdk2-GFP OE (Figure 2D).

**Figure 2:**
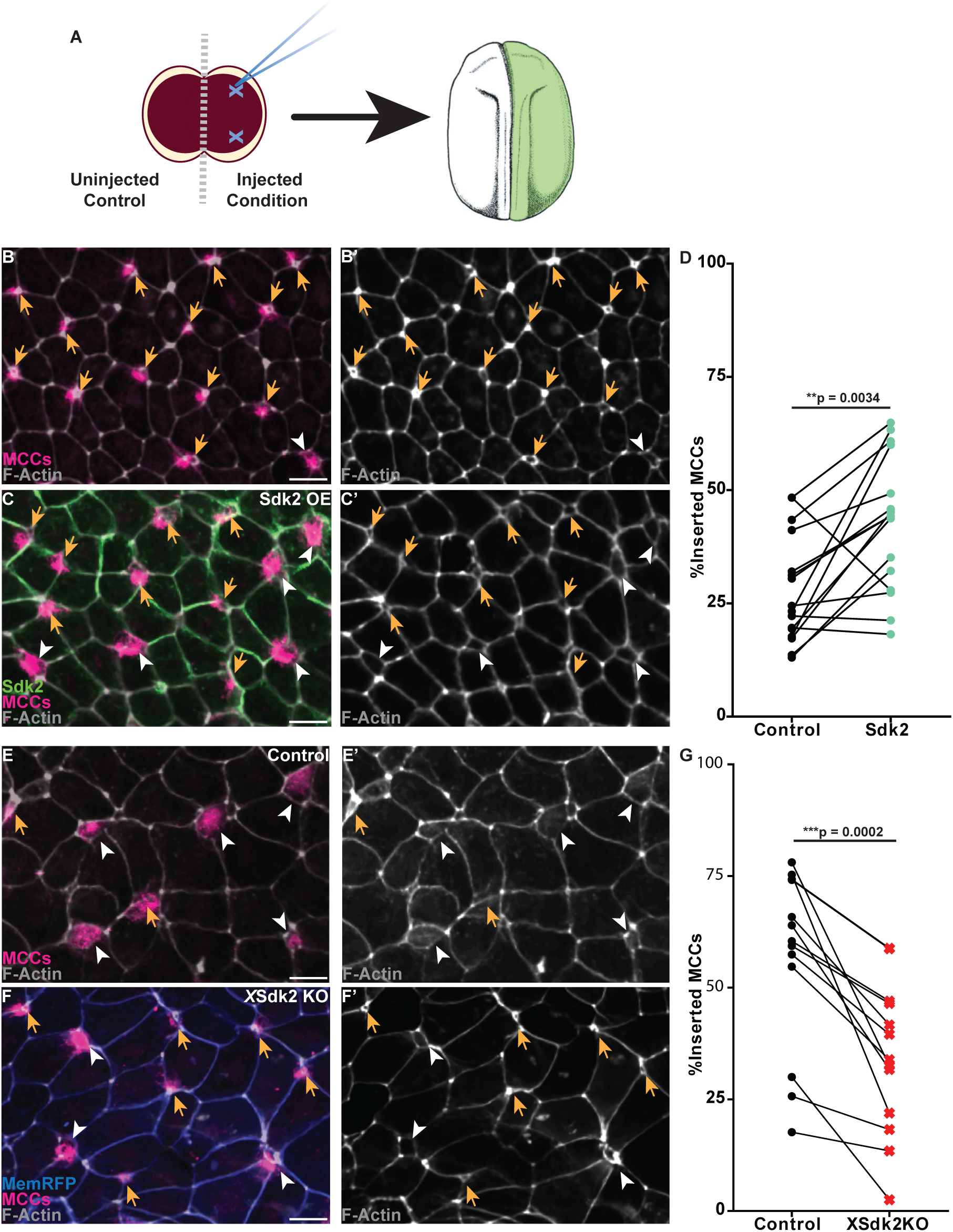
Sidekick2 promotes apical insertion of intercalating Multiciliated cells. (A) Cartoon depiction of mosaic injections with a 2-cell *Xenopus* embryo. Xenopus embryo illustration in A from Nieuwkoop and Faber 1994/Xenbase. (B-C) Z projections of mosaic halves of ST20 embryo marked with MCCs (pink), F-Actin (gray), and Sdk2-GFP (green) (C). (B’-C’) F-Actin alone. D) Quantification of the percent of inserted MCCs from mosaic embryos comparing uninjected and Sdk2-GFP injected sides, n > 50 cells per embryo side from 16 embryos. (E-F) Z projections of mosaic halves of ST21 embryo marked with MCCs (pink), F-Actin (gray), and Membrane-RFP to visualize the Cas9 and guides injected half (blue) (C). (E’-F’) F-Actin alone. G) Quantification of the percent of inserted MCCs from mosaic embryos comparing uninjected and Cas9 / Sdk2 gRNA injected sides, n > 50 cells per embryo side from 13 embryos. (B-C, E-F) Scale bars are 20µm. Yellow arrows indicate MCCs with apical areas < 30μm^2^ and white arrow heads indicate MCCs with apical areas > 30μm^2^. (D and G). Two tailed p-values are calculated with a paired Wilcoxon test.

Next we tested the effects of knocking out (KO) Sdk2. For these experiments, we utilized the CRISPR Cas9 system, designing 4 guides targeting both copies of the *X*Sdk2 gene (Figure S2A). To distinguish the injected embryonic halves, we co-injected a membrane marker (MemRFP) alongside the Cas9 and guides (Figure 1F). Importantly, at ST21 when most MCCs have successfully inserted, we observed a significant delay in MCCs insertion on the crispant half of the embryos compared to the control (Figure 2E-G). This reduction in insertion was not due to injecting Cas9, because mosaic injections with Cas9 and MemRFP alone had no significant effect on MCC insertion (Figure S2B-D). Taken together, these results suggest Sdk2 expression at impacts MCC RI. Importantly, our results suggest that Sdk2 provides an instructive cue at McJs that facilitates insertion, rather than serving as an obstacle to pass through.

### The Intra- and Extracellular Domains Impact Sdk2 Dynamics at Junctions

Epithelial remodeling is facilitated by dynamic changes to junctional proteins^35,36^. We therefore wanted to test the dynamics of Sdk2 at both BcJ and McJs. Additionally, we wanted to explore how the intracellular PDZ binding motif and the extracellular domain of Sdk2 contribute to these dynamics. To target the role of the intracellular domain, we created the mutant sdk2ΔCM-GFP by deleting the last 25 amino acids, including the highly conserved C-terminus, AGFSSV. Studies in *Drosophila* have implicated the highly conserved C-terminal PDZ binding domain as an important protein interaction site, and we wanted to interrogate how removal of this domain may affect Sdk2 junctional dynamics^13,22,24,26,37^. Furthermore, previous reports have suggested multiple roles for the different IG domains and FN-like repeats within the Sdk2 extracellular region related to dimerization and proper protein architecture^24–26^. We therefore generated a construct removing the entire extracellular domain, GFP-sdk2ΔExC. Both mutants localize to McJs; sdk2ΔCM-GFP has decreased intensity and GFP-sdk2ΔExC increased intensity at McJs when compared to Sdk2-GFP (Figure S3A-B).

To test the dynamics of Sdk2 and our mutants, we utilized Fluorescence Recovery After Photobleaching (FRAP). We quantified the dynamics of Sdk2 by photobleaching at McJs (Figure 3B) as well as BcJs (Figure 3F), imaging in 2.1 second increments over 2.5 minutes. A pre-bleach baseline of 5 frames was recorded prior to bleaching. These experiments were performed using ST12 embryos, a simple epithelium that is not confounded by specialized junctional interactions that could occur within the complex epithelium at later STs.

**Figure 3:**
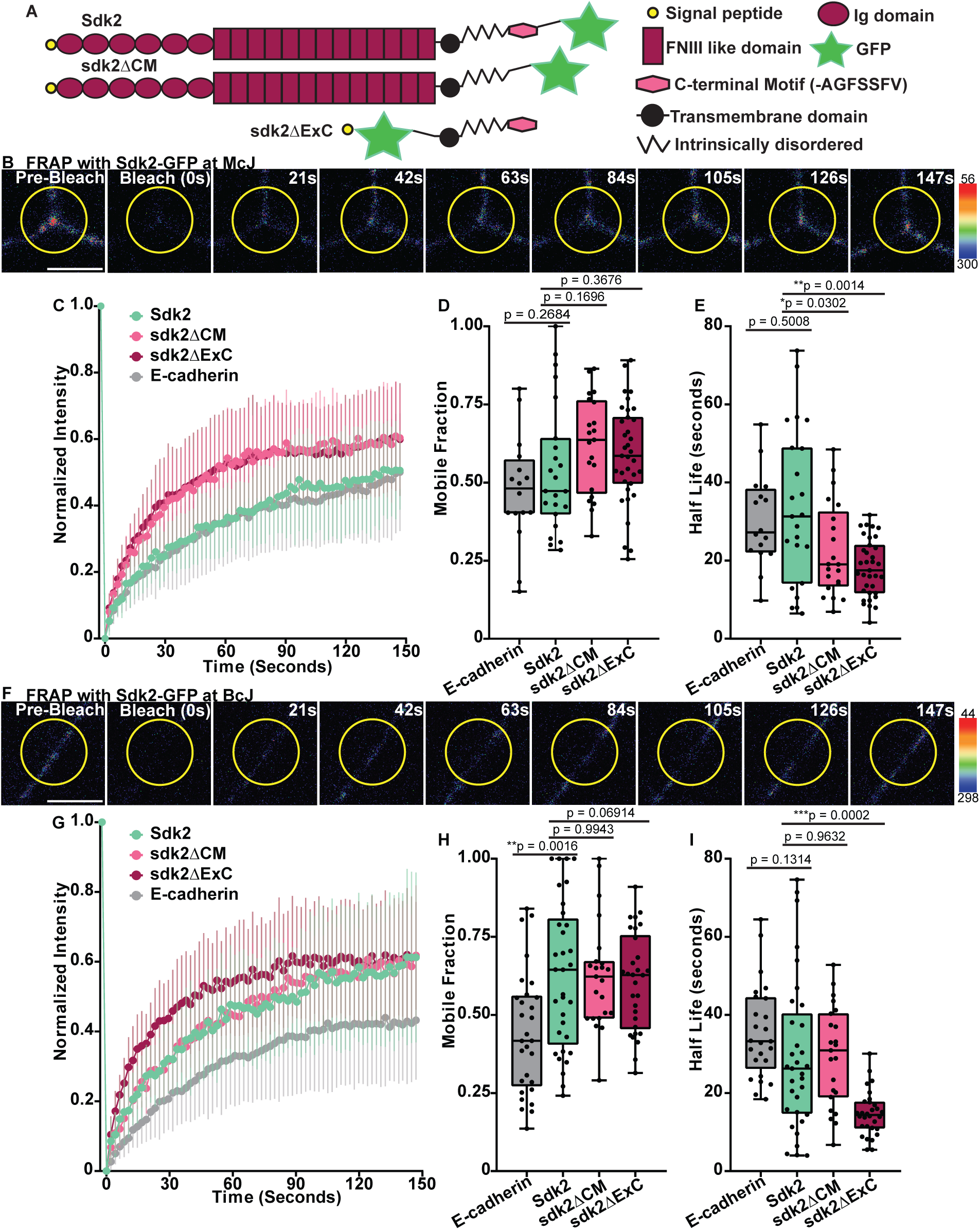
Domain modulation of Sidekick2 junctional dynamics. (A) Cartoon depiction of Sdk2-GFP and domain diderences between the mutants, sdk2ΔCM and sdk2ΔExC. (B) Representative still frames from FRAP analysis with Sdk2-GFP at a McJ. (C-E) fluorescence recovery curves (C), mobile fractions (D) and half-lives (E) for Sdk2-GFP (aquamarine), sdk2ΔCM-GFP (light pink), GFP-sdk2ΔExC (burgundy), and E-cadherin-GFP (gray) at McJ. n = 23 McJ from 7 embryos (Sdk2-GFP); 21 McJs from 6 embryos (sdk2ΔCM-GFP); 35 McJs from 10 embryos (GFP-sdk2ΔExC); 16 McJs from 5 embryos (E.Cadherin-GFP). (F) Representative still frames from FRAP analysis with Sdk2-GFP at a BcJ. (G-I) fluorescence recovery curves (G), mobile fractions (H) and half-lives (I) for Sdk2-GFP (aquamarine), sdk2ΔCM-GFP (light pink), GFP-sdk2ΔExC (burgundy), and E-cadherin-GFP (gray) at BcJ. n = 33 BcJ from 9 embryos (Sdk2-GFP); 23 BcJs from 7 embryos (sdk2ΔCM-GFP); 30 BcJs from 9 embryos (GFP-sdk2ΔExC); 29 BcJs from 7 embryos (E-cadherin-GFP). (B, F) Time points pre- and post-bleaching are shown; time is measured in seconds and the scale bar is 5μm. (D-E, H-I) Two-sided p-values are determined with an unpaired Welch’s T-test.

Upon FRAP with Sdk2-GFP, we observed similar recovery speeds between McJs and BcJs (Figure S3C, E). However, there was an increase in fluorescent recovery (i.e. mobile fracation) at BcJs compared to McJs, suggesting Sdk2 is more stable at McJs than BcJs (Figure S3C, D). To contextualize our FRAP findings, we compared Sdk2 dynamics against E-cadherin. FRAP with either Sdk2-GFP or E-cadherin-GFP at McJs had similar dynamics, with no significant difference in the mobile fractions or half-lives (Figure 3C-E). At BcJs, we observed significantly more fluorescence recovery with Sdk2 than E-cadherin and Sdk2 trending toward a faster recovery (Figure 3G-I). This indicates that Sdk2 is more mobile along bicellular junctions than E-cadherin, which is supported with *Drosophila* studies that show Sdk2 is important for T-1 junctional remodeling during embryonic development^13–15,27^.

FRAP analysis of our mutants at McJs vastly differed from full-length Sdk2 but were similar to each other (Figure 3C-E). Comparison of the mobile fractions indicates that sdk2ΔCM-GFP has significantly more fluorescence recovery compared to Sdk2-GFP (Figure 3C-D). Both mutants recovered significantly faster than the full-length protein (Figure 3E). In contrast, at BcJs, the two mutants had fluorescence recovery comparable to Sdk2-GFP, but GFP-sdk2ΔExC did recover significantly faster than Sdk2-GFP or sdk2ΔCM-GFP (Figure 3G-I). Comparison of each mutant itself at McJ and BcJ displayed similar recoveries, though with alterations in recovery speed (Figure S3F-K). These results indicate that both the intracellular and extracellular domains have roles in stabilizing Sdk2 at McJs (Figure 3C).

### Sdk2 intracellular domain influences the timing of Multiciliated Cell Radial Intercalation

Given the FRAP differences seen between Sdk2-GFP with sdk2ΔCM-GFP and GFP-sdk2ΔExC at McJs, we were curious how these altered dynamics would affect MCC RI. Mosaic injections with sdk2ΔCM-GFP showed no significant changes with MCC RI (Figure 4A-C), indicating that this construct does not function as a dominant negative (DN) by interfering with the endogenous protein. In contrast, mosaic injection of GFP-sdk2ΔExC caused a significant delay in MCC RI, with an observable amount of MCCs still beneath the outer epithelium (52.45% with GFP-sdk2ΔExC vs 39.07% with the control), not yet penetrating (Figure 4D-F). This result stresses the importance of the extracellular domain in the increase in MCC RI observed with OE of full-length Sdk2-GFP (Figure 2B-D) but also suggests that the intracellular domain can act as a DN, presumably by creating non-productive intracellular interactions that interfere with endogenous Sdk2 function.

**Figure 4:**
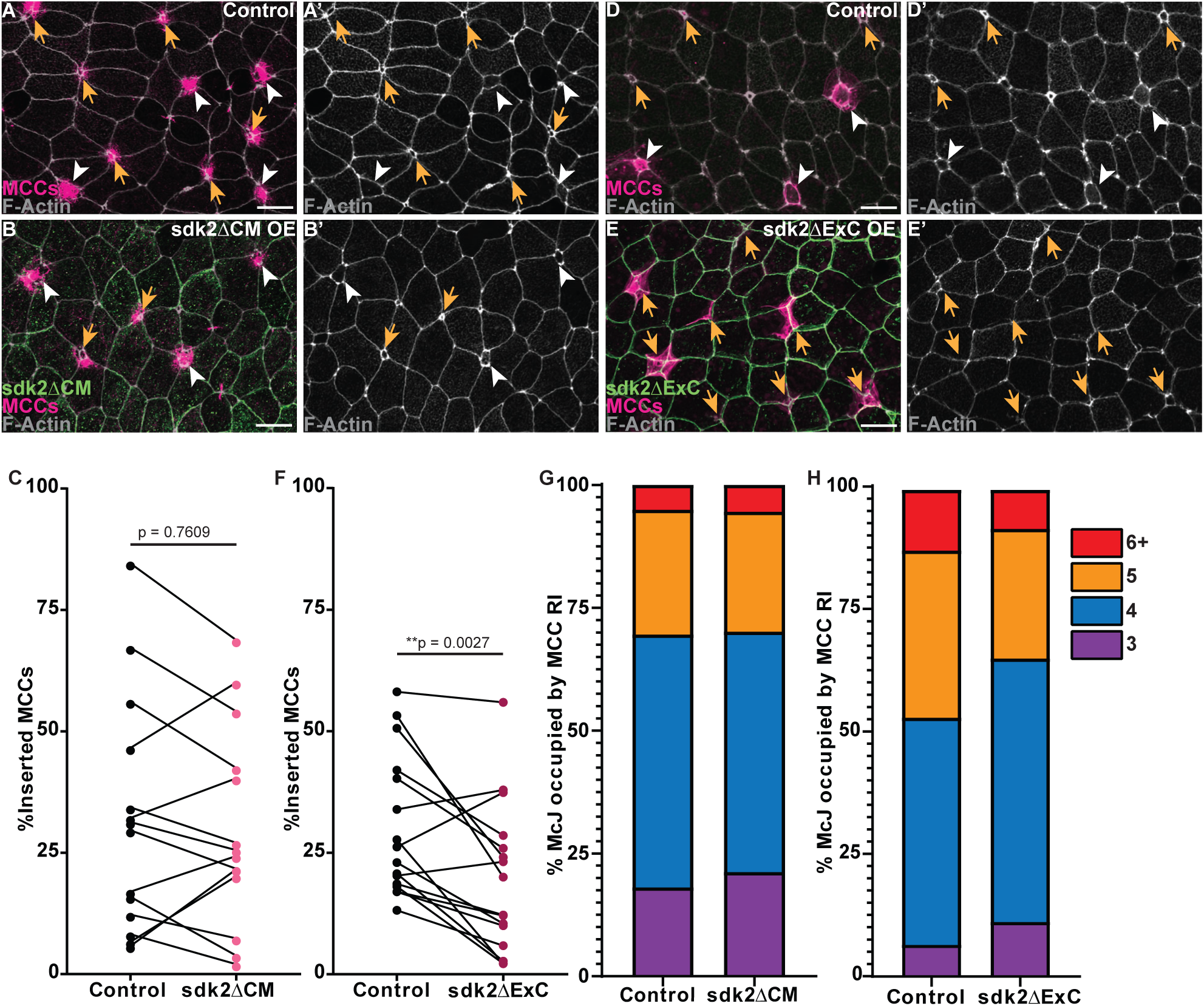
Sidekick2 intracellular domain modulates the timing and vertex preference of Multiciliated cell intercalation. (A-B) Z projections of mosaic halves of ST20 embryo marked with MCCs (pink), F-Actin (gray), and sdk2ΔCM-GFP (green) (C). (B’-C’) F-Actin alone. C) Quantification of the percent of inserted MCCs from mosaic embryos comparing uninjected and sdk2ΔCM-GFP injected sides, n > 50 cells per side from 14 embryos. (D-E) Z projections of mosaic halves of ST20 embryo marked with MCCs (pink), F-Actin (gray), and GFP-sdk2ΔExC (green) (C). (B’-C’) F-Actin alone. (F) Quantification of the percent of inserted MCCs from mosaic embryos comparing uninjected and GFP-sdk2ΔExC injected sides, n > 50 cells per side from 16 embryos. (G-H) percentage of apically inserted MCCs (apical areas > 30 μm ^2^) occupying a McJ composed of either 3, 4, 5, or 6+ cells of mosaic embryos with either sdk2ΔCM-GFP (G) or GFP-sdk2ΔExC (H). Chi-squared analysis of McJ occupancy with sdk2ΔCM-GFP is p = 0.7116 (G) and with GFP-sdk2ΔExC is p <0.0001**** (H). (A-B, D-E) Scale bars are 20μm. Yellow arrows indicate MCCs with apical areas < 30μm^2^ and white arrow heads indicate MCCs with apical areas > 30μm^2^. Two-tailed p-values are calculated with a paired Wilcoxon test.

MCCs typically insert faster at higher order vertices (5-6) compared to lower order vertices (3-4)^29,34^. Importantly, as MCCs probe McJs they form homophilic attachments with the extracellular domain of LSR, and during this attachment remodel the epithelium by merging junctions to create higher order vertices (5-6+ groupings of cells opposed to 3-4)^29^. We surveyed the order of vertices where MCCs intercalate, scoring groupings of 3, 4, 5 or 6+ (Figure S4A). With the OE of Sdk2-GFP, *X*Sdk2 KO, or sdk2ΔCM-GFP we saw no significant change in the magnitude of McJs intercaltion sites (Figure S4B-C and Figure 4G). However, OE of GFP-sdk2ΔExC showed a significant shift of MCC insertion at lower order vertices (3-4) compared to control (Figure 4H). This shift towards lower, more restrictive, vertices could underlie the delay in MCC insertion since these are known to insert more slowly^34^.

## Discussion

Here we report that Sdk2-GFP localizes to epithelial AJ vertices in the vertebrate embryonic *Xenopus* model system (Figure 1). Using the developmental process of MCC RI, we show that Sdk2 affects MCC penetrance at epithelial vertices (Figure 2), and that this process hinges on the Sdk2 intracellular domain (Figure 4D-F). Although, our FRAP analysis indicates that both the extracellular and intracellular domains contribute to Sdk2 dynamics at vertices (Figure 3).

Our findings demonstrate that MCCs require Sdk2 for the proper timing of RI in *Xenopus* embryonic development, with mosaic OE of Sdk2-GFP increasing MCC RI and mosaic Crispant KO showing a delay (Figure 2B-G). The OE of WT Sdk2 facilitating RI suggests that it provides an instructive extracellular cue to intercalating cells. However, that extracellular cue alone is not sufficient as the OE of sdk2ΔCM failed to facilitate RI (Figure 4A-C). In vitro studies have shown that homophilic adhesion is sufficient with the extracellular domain alone^24^, so we do not anticipate our sdk2ΔCM-GFP construct is unable to adhere. The combination of these results implicates the intracellular domain as an important mediator of Sdk2 function. Consistent with this, OE of sdk2ΔExC, acts as a DN overpowering the endogenous protein, potentially via non-productive intracellular interactions (Figure 4F). Similarly, a recent paper in *Drosophila* has shown that the deletion of the last 5 amino acids impacting the PDZ binding motif was sufficient to disrupt trachea development^26^.

While studies in *Drosophila* have shown an interaction between Sdk and the *Drosophila* ZO-1 homologue, Pyd^13,26^, we observed no colocalization of these proteins (Figure S1D-E). Those same studies demonstrated that Sdk interacts with Cno^13,26^, the homologue of AJ scaffolding protein Afadin. A current preprint with vertebrate epithelium shows that Afadin recruits the kinase PAK4 to junctions and that this kinase activity at junctions is important for junctional remodeling^38^. Intriguingly, a recent *Drosophila* study shows Sdk phosphorylation by Mbt, the PAK4 homologue, is important for Sdk localization to vertices^26^. While future studies are required to determine if there is an interaction between Sdk2, Afadin, and PAK4, it is tempting to speculate that this could be a mechanism for junctional remodeling and establishing new epithelial vertices.

Collectively, our findings suggest Sdk2 is a specialized McAJ protein that regulates epithelial integrity and facilitates RI. Future studies are required to determine if Sdk2 broadly expressed across vertebrate epithelium. We anticipate that the growing appreciation of the importance of McJ in epithelial regulation will provide novel insights for understanding development and disease.

## Acknowledgements

We are grateful to Ann Miller, University of Michigan, for providing us the *Xenopus* LSR antibody. This work was supported with R01GM113922 grant from NIH-NIGMS to BM. This work was funded by the JSPS (Japan Society for the Promotion of Science) KAKENHI (Grant Number 24K09456), 2024 Narishige Zoological Science Award, and the 56th Research Grant from the Naito Foundation to T.H. We would like to thank the National Xenopus Resource center and Xenbase for their important community service that were critical for this work.

## Materials and Methods

### Experimental model details

Samples were generated with *Xenopus laevis* in vitro fertilizations using previous established standards by the Northwestern University Institutional Animal Care and Use Committee (# IS00001612). Wild type (WT) and Tub-mRFP (tuba1a:MyrPalm-mRFP1, NRX_0074; for marking MCCs) transgenic lines were obtained from Xenopus1 (WT) or the National Xenopus Resource Center (NXR; WT and TubA1A). Embryos in this study were used early in development, prior to sexual differentiation, so sex cannot be used as a biological variable.

### Injections

Wild-type or transgenic embryos were injected at either the two-, or four-cell stages. All mosaic injections were conducted at the two-cell stage to ensure right-left patterning mosaicism (as illustrated in Figure 2A^39^). Embryos for mosaic OE of Sdk2-GFP, sdk2ΔCM-GFP, and GFP-sdk2ΔExC (Figure 2 and Figure 4) were injected with 50pg of mRNA. Embryos were injected with 15-20pg of Sdk2-GFP and coinjected with either 40pg Tricellulin-RFP, 15pg Occludin-mCherry, 20pg E-cadherin-2xmCherry, 50pg ZO-1-mCherry (Figure 1, Figure S1) or 15pg Mem-RFP (Figure 3). All injected mRNAs were synthesized with the Sp6 mMessage Machine kit (Life Technologies, AM1340) and purified by RNeasy MiniElute Cleanup Kit (QIAGEN, 74204).

### CRISPR

Embryos were injected at the the two-cell stage to ensure mosaicism with 250pg of Cas9 (PNA Bio Inc) and 25pg MemRFP as a control or with 250pg of Cas9, 250pg of gRNAs (in total, 62.5pf of each guide) (Synthego Inc), and 25pg MemRFP to KO *X*Sdk2. gRNAs used were designed using CHOPCHOP (Labun, et al.^40^). Four guides were used as Sdk2 is a pseudo tetraploid gene in *Xenopus Laevis*. Sequences of the guides used are: Chr9_10L Sdk2 Exon1 (- strand, CTGCTAGTCCGTGCAGTGCTGGG), Chr9_10L Sdk2 Exon3 (+ strand, TCTCTTGACCGGTCTCATGCTGG), Chr9_10S Sdk2 Exon6 (- strand, TCTCTGGACAGGACTCACGCTGG), Chr9_10S Sdk2 Exon1 (+ strand, TCTCTGGACAGGACTCACGCTGG).

### Immunofluorescence

Most samples were fixed with 4% PFA/PBS. E-cadherin (DSHB 5D3, [1:200]) and ZO-1(Invitrogen #61-7300, [1:200]) stained samples were fixed with Dent’s (80% methanol and 20% DMSO, 24 hours at -20°C), with the injected Sdk2 in these samples marked with anti-GFP (either Invitrogen A-11120 or ABCAM ab13970). Post-fixation, embryos were blocked in 10% Heat Inactivated Goat Serum (HIGS) for an hour, with primary and secondary antibodies diluted in 5% HIGS with overnight incubations. To visualize MCCs, embryos were either generated with transgenic Tub-mRFP lines or stained for acetylated α-tubulin (T7451; Sigma-Aldrich, [1:500]). LSR (custom anti-body produced in rabit, Higashi et al.^41^ [1:50]). Cy-2, Cy-3, or Cy-5 conjugated goat anti-mouse or anti-rabbit (Thermo Fisher Scientific) were used at 1:750 dilutions. Cell borders were marked by staining for F-Actin with either Acti-Stain Phalloidin 670 (#PHDN1, [1:500]) or Alexafluro Plus 405 Phalloidin (Invitrogen A30104, [1:400]) diluted in PBST and incubated for 1 hour. Embryos were mounted between coverslips with Fluoro-Gel (Electron Microscopy Sciences).

### Microscopy

Microscopy was performed with a laser-scanning confocal microscope (A1R; Nikon) using a 60 × oil Plan-Apochromat objective lens (NA 1.4) or a Plan Fluor 40× Oil DIC H N2 (NA 1.3). Fixed Embryos were mounted between coverslips with Fluoro-Gel (Electron Microscopy Sciences). Live embryos were mounted in a glass bottom petri dish (MatTek) with a coverslip attached to the slide with vacuum grease. Images with fixed samples multiple x planes are visualized in 0.3-0.5 μm (5-10 μm depth) of either a 512x512 or 1024x1024 pixel area with a pinhole size of 35.76 AU and the suitable laser (450-nm, 525nm, 595nm, and 700nm). Live imaging for Frap experiments were conducted with time-lapse movies scanning of a 1024 x 512 pixel area with a pinhole size of 60.03 AU and a 488-nm laser. Photobleaching was performed in NIS with a 30% laser power for 2.1sec, with a circular ROI that encapsulated a tricellular (3 cell junctions specifically were used) or a square ROI at the middle of bicellular junctions. 5 frames were taken prior to bleaching. Nikon Elements Software was used for all acquisition and image processing.

### Quantification & Analysis

#### Apical Insertion & MCC Vertex Analysis

Z-projections were created to identify inserting MCCs at differing depths in the epithelium. The apical area of intercalating cells was measured by outlining the area of the apical surface marked by F-Actin staining. For analysis, an apical area of 30μm^2^ was set as a threshold of a successful MCC insertion. The percentage of MCC successful insertions indicates MCCs > 30μm^2^/total MCCs counted, cells with an indeterminable apical area (∼<3 μm^2^) are included in the analysis and were assigned an apical area of 0μm^2^. Alongside apical size, the magnitude of cells composing a McJ where MCCs intercalated at was recorded an grouped into one of four categories: 3, 4, 5, or 6+ cells. As categorical data this was analyzed with a Chi-square test to determine statistical significance, but is presented as percentages of all cells counted to compare raw amounts.

#### Quantification of FRAP recovery

For both McJ and BcJ FRAP sets, the intensity of the bleached region of interest, a reference junction, and a non-junction background area were quantified in all frames using a circular ROI (2.82 μm diameter). In both sets a junction from an opposing cell (ie not another junction shared between the same cell being bleached) was chosen for the control. All McJ bleached were composed of only 3 cells. The ROIs were measured from aligned images with NIS Elements to correct for drift. These measurements were organized with Microsoft Excel where both the bleached and reference ROIs were corrected by subtracting the background/autofluorescence ROI. We normalized and constrained our values using the methods from van den Goor et al.^42^ The intensities were normalized with the formula:

I_norm_ (t) = (I _ref pre_/I_ref_ (t))* (I_frap_(t)/I_frap pre_). The normalized Intensities were constrained from 1 to 0 with the formula: I_norm1_ (t) = (I_norm_(t) – I_norm_ (t_bleach_)) / (I _norm pre_ – I _norm_ (t _bleach_)). The I_frap pre_ value was the average intensity of the 5 frames taken prior to bleaching. Each recovery curve was fit to a exponential one phase association equation in GraphPadPrism, with the constraints that Y_0_ = 0 and Plateau had to be between 0 and 1, to determine the individual mobile fractions and half-lives. Data represented in the frap curves are the compilation of three biological replicates for each condition.

#### Quantification of Fluorescent Intensity at McJ

Using the first frame (pre bleach) of a selection of our FRAP videos, we measured a circular ROI (1.8 μm diameter) centered at every tricellular junction in the field of view. 6 embryos were measured (2 from each of the 3 biological replicates), with 2-3 images from each embryo. Only tricellular junctions (groups of three cells) were measured. These intensities were corrected by subtracting background/autofluorescence with an ROI not at junctions.

### Image preparation & Statistical Analysis

Some images were smoothed and processed for figure presentation only. Raw images were used for all analyses. Statistical significance for MCC RI was performed with a Wilcoxon matched-pairs signed rank test, Mobile-Fraction and Half-Life were performed with unpaired t test with Welch’s correction, MCC RI at different orders of McJs was performed with a Chi-Square test, and differences of fluorescence intensity at McJ between Sdk2-GFP and mutants was performed with a Mann-Whitney test. For all statistical analyses, ^∗^p < 0.05, ^∗∗^p < 0.01, and ^∗∗∗^p < 0.001. Images were analyzed in NIS Elements, recorded measurements were organized with excel spreadsheets and graphs and statistics were calculated with Graph Pad Prism.

