## Supplemental Data for "Sidekick2 facilitates multiciliated cell penetration through multicellular adherens junctions"

**Figure S1:** (A-E) Z projections of embryos marked with MsSdk2-GFP (green) F-Actin (gray) together with Tricellulin-RFP at ST21 (A), Occludin-mcherry at ST20 (B), E-cadherin antibody at ST21 (C), ZO-1-mCherry at ST21 (D), and ZO-1 antibody at ST19 (E) (pink). Scale bars are 20μm. (A'-E') Side projections from boxed regions in (A-E).

**Figure S2:** (A) Sequences of the four CRISPR guides used to target XSdk2 located on the tetraploid chromosome 9_10. (B-C) Z projections of mosaic halves of ST21 embryo marked with MCCs (pink), F-Actin (gray), and Membrane-RFP to visualize the Cas9-injected half (blue) (C). (B'-C') F-Actin alone. Scale bars are 20μm. Yellow arrows indicate MCCs with apical areas < 30$\mu m^2$ and white arrow heads indicate MCCs with apical areas > 30$\mu m^2$. D) Quantification of the percent of inserted MCCs from mosaic embryos, n > 50 cells per embryo side from 15 embryos. Two-tailed p-values are calculated with a paired Wilcoxon test.

**Figure S3:** (A) Representative images showing expression of Sdk2-GFP, sdk2ΔCM-GFP, and GFP-sdk2ΔExC at junctions. Scale bars are 20μm. (B)Mean fluorescent intensities at McJs with Sdk2-GFP (aquamarine, n = 154 McJs), sdk2ΔCM-GFP (light pink, n = 132 McJs), and GFP-sdk2ΔExC (burgundy, n = 140 McJs). Counts were taken from 6 embryos for each condition. (C-E) fluorescence recovery curves with Sdk2-GFP (C), sdk2ΔCM-GFP (D), and GFP-sdk2ΔExC (E) at McJs (orange) and BcJs (light blue). (F,H, J) Mobile fractions and (G, I, K) half-lives of Sdk2-GFP (F-G), sdk2ΔCM-GFP (H-I), and GFP-sdk2ΔExC (J-K) at McJs (orange) and BcJs (light blue). Two-tailed p-values are calculated with a Mann-Whitney (B) or a paired Wilcoxon test (F-K).

**Figure S4:** (A) Representative images showing MCC intercalation at McJs composed of 3, 4, 5, or 6 cells. Scale bars are 5μm. (B-C) percentage of apically inserted MCCs (apical areas > 30 $\mu m^2$) occupying a McJ composed of either 3, 4, 5, or 6+ cells of mosaic embryos with either *X*sdk2 KO (B) or Sdk2-GFP (C). Chi-squared analysis of McJ occupancy with sdk2ΔCM-GFP is p = 0.1841 (B) and with Sdk2-GFP is p = 0.5435 (C).

#### Figure S1.

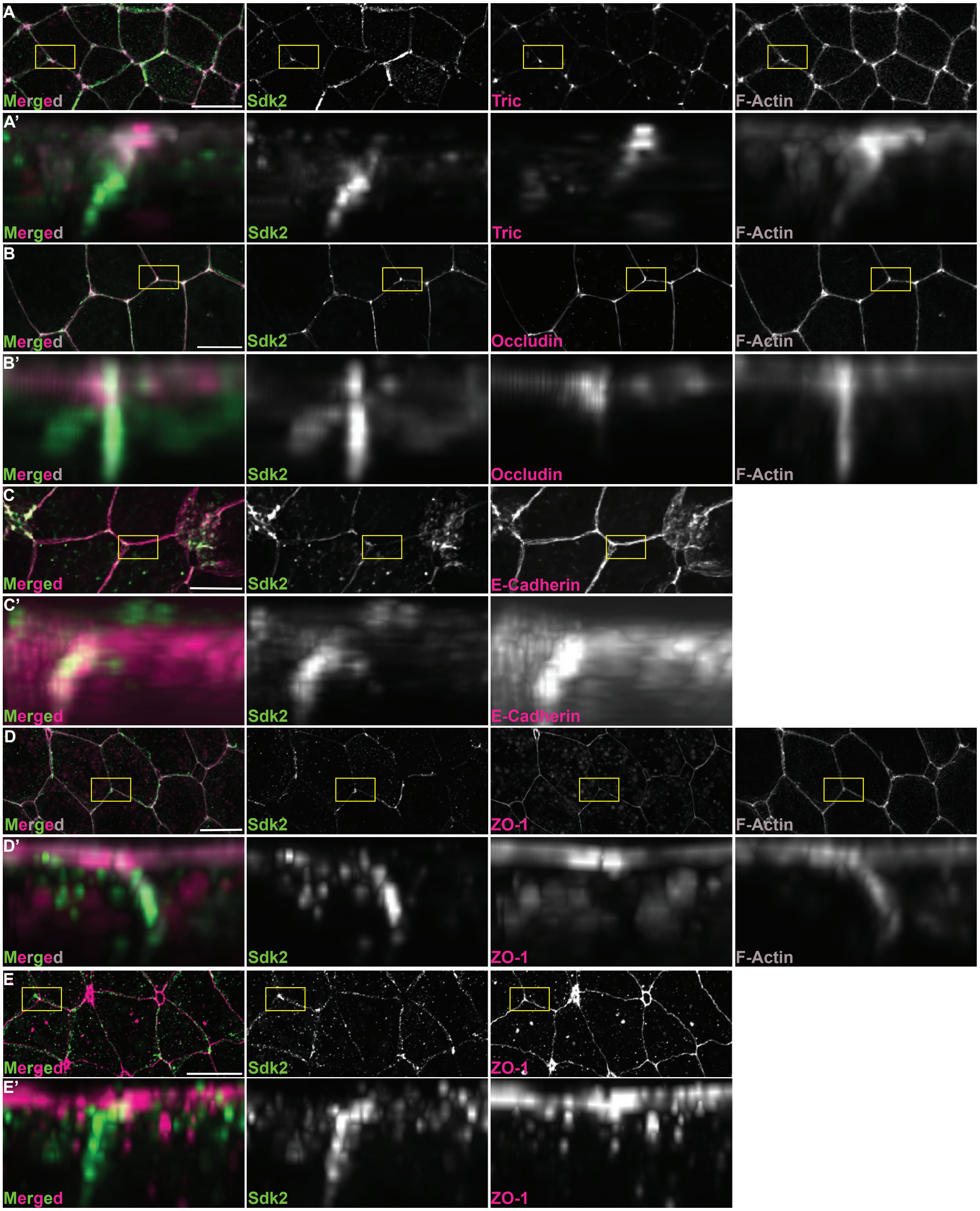

A
Merged
Sdk2
Tric
F-Actin
A'
Merged
Sdk2
Tric
F-Actin
B
Merged
Sdk2
Occludin
F-Actin
B'
Merged
Sdk2
Occludin
F-Actin
C
Merged
Sdk2
E-Cadherin
C'
Merged
Sdk2
E-Cadherin
D
Merged
Sdk2
ZO-1
F-Actin
D'
Merged
Sdk2
ZO-1
F-Actin
E
Merged
Sdk2
ZO-1
E'
Merged
Sdk2
ZO-1

#### Figure S2.

A
**Chr9_10L:**
**(+)TCTCTTGACCGGTCTCATGCTGG**
**(-)CTGCTAGTCCGTGCAGTGCTGGG**

**Chr9_10S:**
**(+)TCTCTGGACAGGACTCACGCTGG**
**(-)CACCACGCTTGTATTGTGCGGGG**

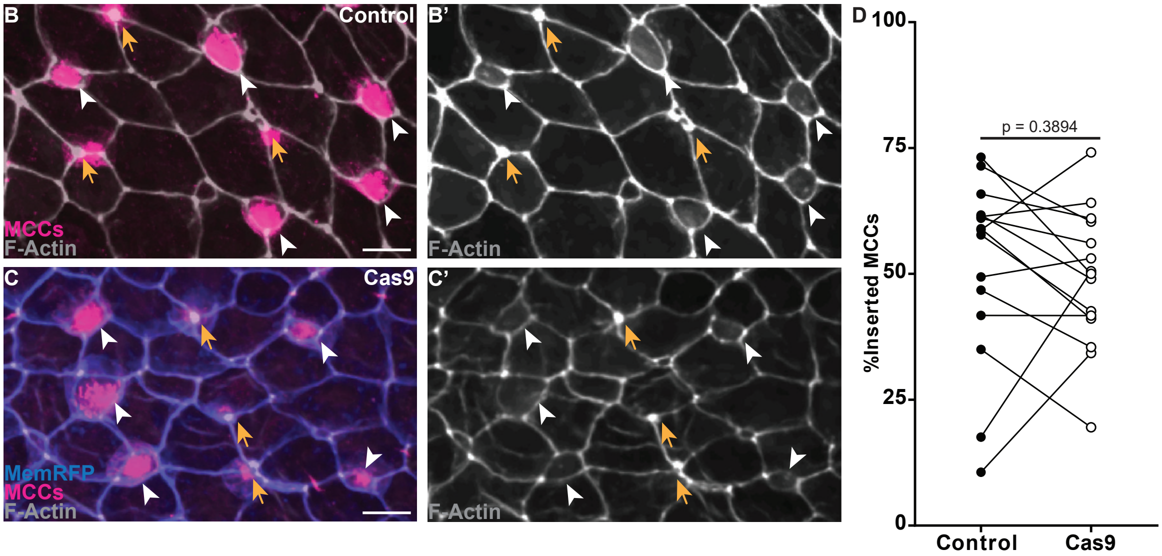

### Figure S3.

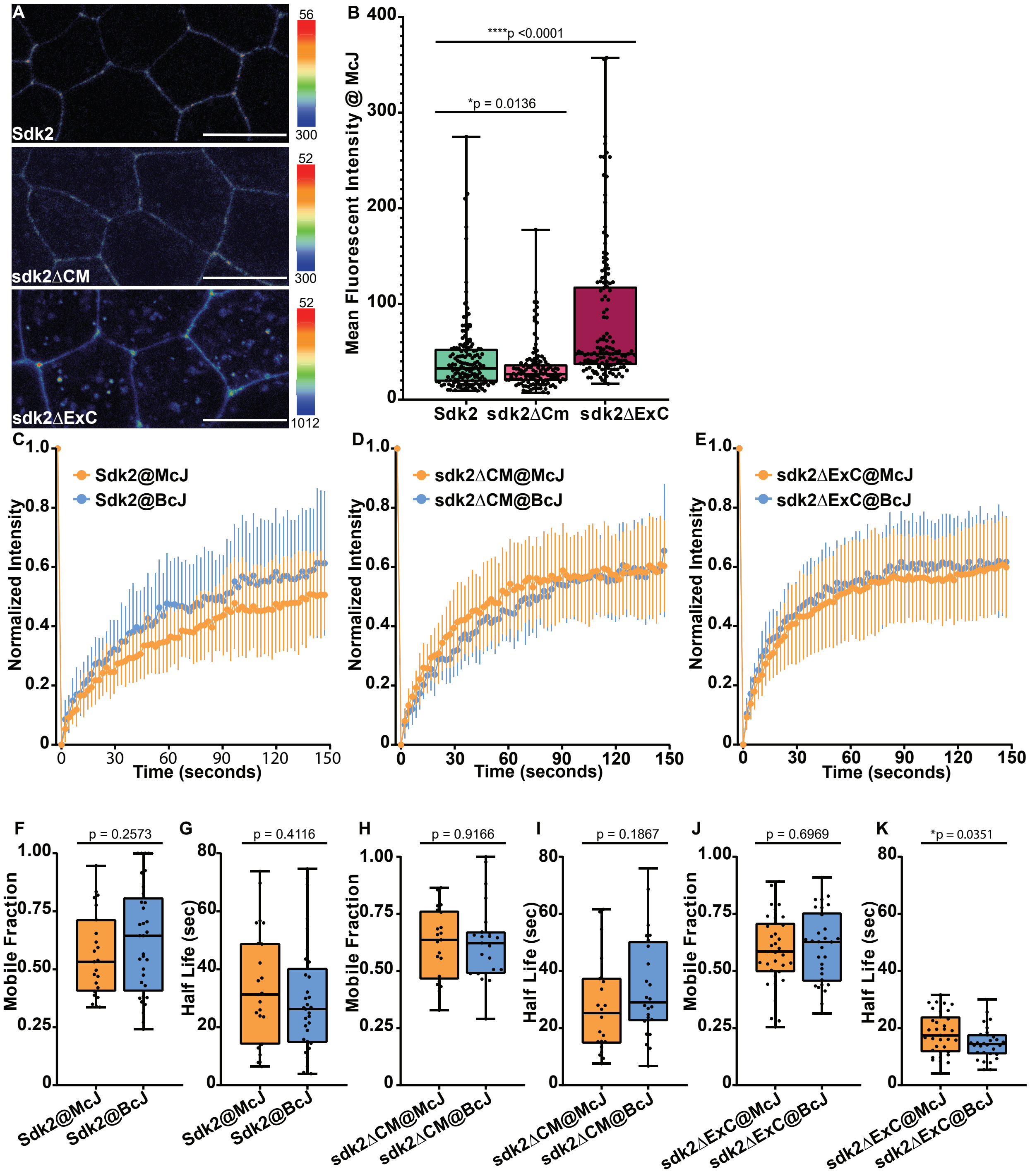
A
Sdk2
sdk2ΔCM
sdk2ΔExC
56
300
52
300
52
1012
B
Mean Fluorescent Intensity @ McJ
****p <0.0001
*p = 0.0136
Sdk2
sdk2ΔCm
sdk2ΔExC
C
D
E
Sdk2@McJ
Sdk2@BcJ
sdk2ΔCM@McJ
sdk2ΔCM@BcJ
sdk2ΔExC@McJ
sdk2ΔExC@BcJ
Normalized Intensity
Time (seconds)
F
G
H
I
J
K
p = 0.2573
p = 0.4116
p = 0.9166
p = 0.1867
p = 0.6969
*p = 0.0351
Mobile Fraction
Half Life (sec)

#### Figure S4.

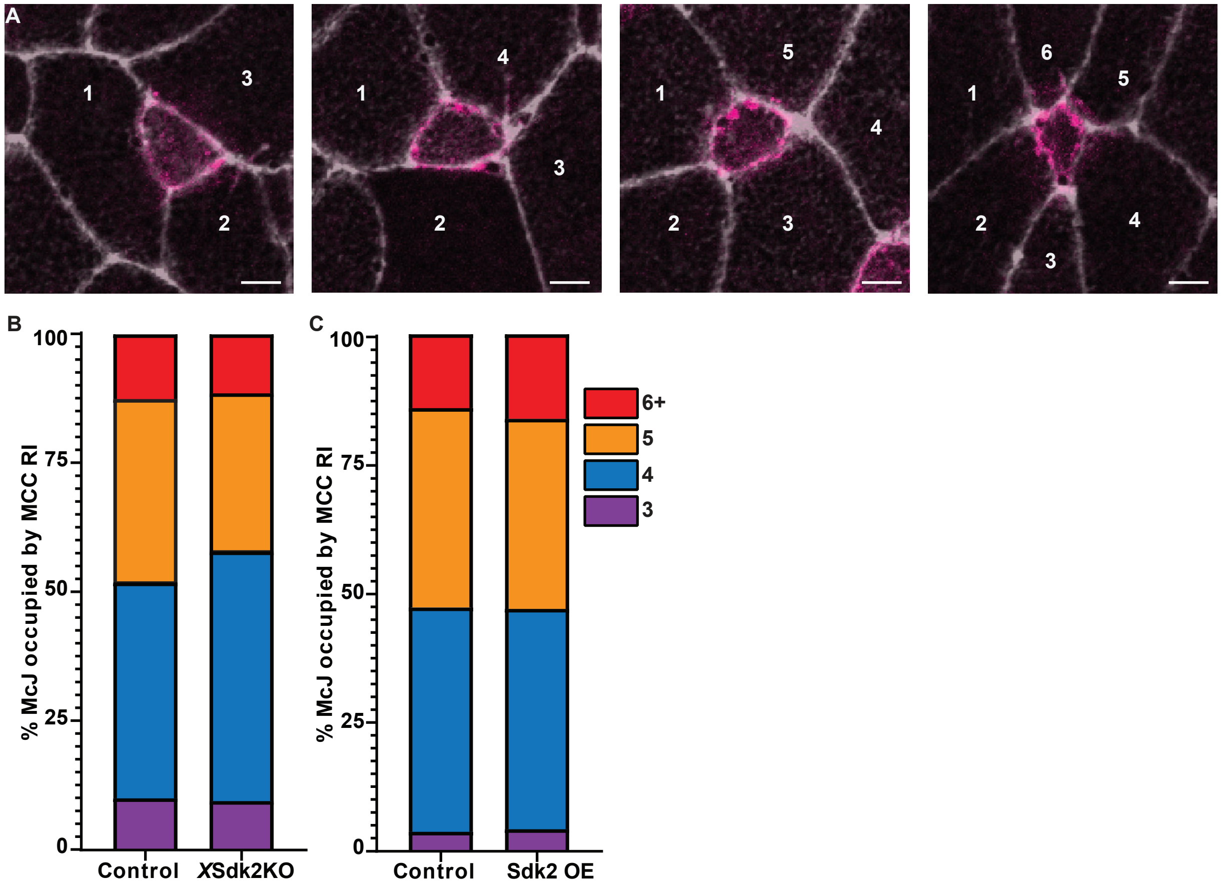
A
1
2
3
4
5
6
B
C
100
75
50
25
0
% McJ occupied by MCC RI
Control
XSdk2KO
Sdk2 OE
6+
5
4
3
